# Size-Dependent Secretory Protein Reflux into the Cytosol in Association with Acute Endoplasmic Reticulum Stress

**DOI:** 10.1101/573428

**Authors:** Patrick Lajoie, Erik L. Snapp

## Abstract

Once secretory proteins have been targeted to the endoplasmic reticulum (ER), the proteins typically remain partitioned from the cytosol. If the secretory proteins misfold, they can be unfolded and retrotranslocated into the cytosol for destruction by the proteasome by ER-associated protein Degradation (ERAD). Here, we report that correctly folded and targeted luminal ER fluorescent protein reporters accumulate in the cytosol during acute misfolded secretory protein stress in yeast. Photoactivation fluorescence microscopy experiments reveal that luminal reporters already localized to the ER relocalize to the cytosol, even in the absence of essential ERAD machinery. We named this process “ER reflux.” Reflux appears to be regulated in a size-dependent manner for reporters. Interestingly, prior heat shock stress also prevents ER stress-induced reflux. Together, our findings establish a new ER stress-regulated pathway for relocalization of small luminal secretory proteins into the cytosol, distinct from the ERAD and pre-emptive quality control pathways.

## INTRODUCTION

The standard model of secretory protein localization in eukaryotes holds that proteins are translated in the cytosol and then trafficked to and inserted into the endoplasmic reticulum (ER) membrane or translocated into the ER lumen in co- and post-translational processes (1, 2). Partitioning of secretory proteins from the cytosol into the ER ensures that secretory proteins fold in the unique ER environment, interact with other partner secretory proteins, and, if appropriate, get secreted out of the ER to other organelles of the secretory pathway or into the extracellular milieu. Secretory proteins that fail to correctly fold can be retrotranslocated from the ER lumen or ER membrane back into the cytosol followed by proteasome mediated destruction in a process termed ER associated degradation or (ERAD) (3-7). Furthermore, some secretory proteins fail to enter the ER due to inefficiencies in the targeting process or sequestration of translocation factors (8-12).

More recently, the Walter group described a condition in which yeast cells impaired in their ability to downregulate the Unfolded Protein Response (UPR) exhibit partial cytosolic localization of a small non-native ER reporter protein, eroGFP (13). The basis of this phenotype is unclear, though the Walter group suggested the cytosolic pool of protein might result from a translocation defect potentially unique to eroGFP, as they reported no altered localization for the resident luminal ER chaperone Kar2p under comparable conditions (13. As our lab performs live cell imaging studies of fluorescent protein-tagged reporters and faithful interrogation of the ER requires that the reporter robustly localizes to the ER, we sought to better understand this phenomenon. Using several reporters with different types of signal peptides, sequences, and sizes, we found that the stress-induced cytosolic pool of accumulation of sfGFP and other small fluorescent protein (FP) reporters was unrelated to translocation defects. Instead, we uncovered a novel pathway, distinct from ERAD, for the movement of correctly folded soluble ER proteins back to the cytoplasm.

## RESULTS

### Acute ER stress leads to accumulation of ER-GFP in the cytosol

Our lab has a long standing interest in environmental reporters in the ER. A fundamental requirement for ER reporters and biosensors is that the proteins must actually localize inside the ER lumen to accurately report on the ER luminal environment. Localization to the ER lumen is achieved typically by attaching targeting (signal peptide) and retrieval/retention (-HDEL) sequences to the reporter. Curiously, we came across a report that described a luminal ER reporter that could be found in the cytosol under ER stress-related conditions. In the Rubio et al. study (13), significant cytosolic localization of a redox reporter protein eroGFP (which contained canonical ER-targeting and retrieval motifs) (14) was observed in mutant yeast cells with a mutated Ire1 (*ire1(D797N,K799N*). Ire1 is the misfolded ER protein sensor and activator of the ER stress pathway, the Unfolded Protein Response (UPR) (15). The Ire1 mutant exhibited impaired attenuation of kinase activity following removal of a misfolded secretory protein stress (DTT). Relevant to our study, cytosol-localized eroGFP was also observed in the absence of application of DTT, which led to the suggestion that these cells had a constitutive translocation defect for secretory proteins. In the following study, we sought to better characterize this phenotype, determine whether it occurred for other ER stresses and reporters, and then attempted to modify ER reporters to improve their ER retention and localization during ER stress.

Our first goal was to determine whether ER misfolded protein stresses led to cytosolic localization of FP reporters. In Fig. 1 A, we observed that sfGFP targeted to the ER with a Kar2 signal sequence and COOH-terminal HDEL ER localization motif (16) localized robustly to the nuclear envelope and peripheral ER in unstressed yeast. Acute treatment with DTT led to accumulation of ER-sfGFP in the cytosol within 30 min of treatment and cytosolic localization became more pronounced with longer treatment times. Similar results were observed with tunicamycin (Tm) a disruptor of GlcNAc phosphotransferase that impairs N-linked glycosylation of secretory proteins. Tm treatment took longer to exhibit the cytosolic accumulation and this is consistent with a delay in impaired N-linked glycosylation due to the need to first deplete existing stores of dolichol-bound N-acetylglucosamine derivatives. Cytosolic accumulation was also observed with an FP with a distinct primary sequence and spectral profile, mCherry (17) (Fig. 1 B). Our results suggested that visible accumulation of cytosolic ER-sfGFP depends on acute ER-stress. In fact, cytosolic accumulation does not require the UPR pathway. Yeast deleted for either of the two key UPR effectors, Ire1 or Hac1, still accumulate cytosolic ER-sfGFP with acute misfolded secretory protein stress (Fig. 1 C).

**Figure 1.**
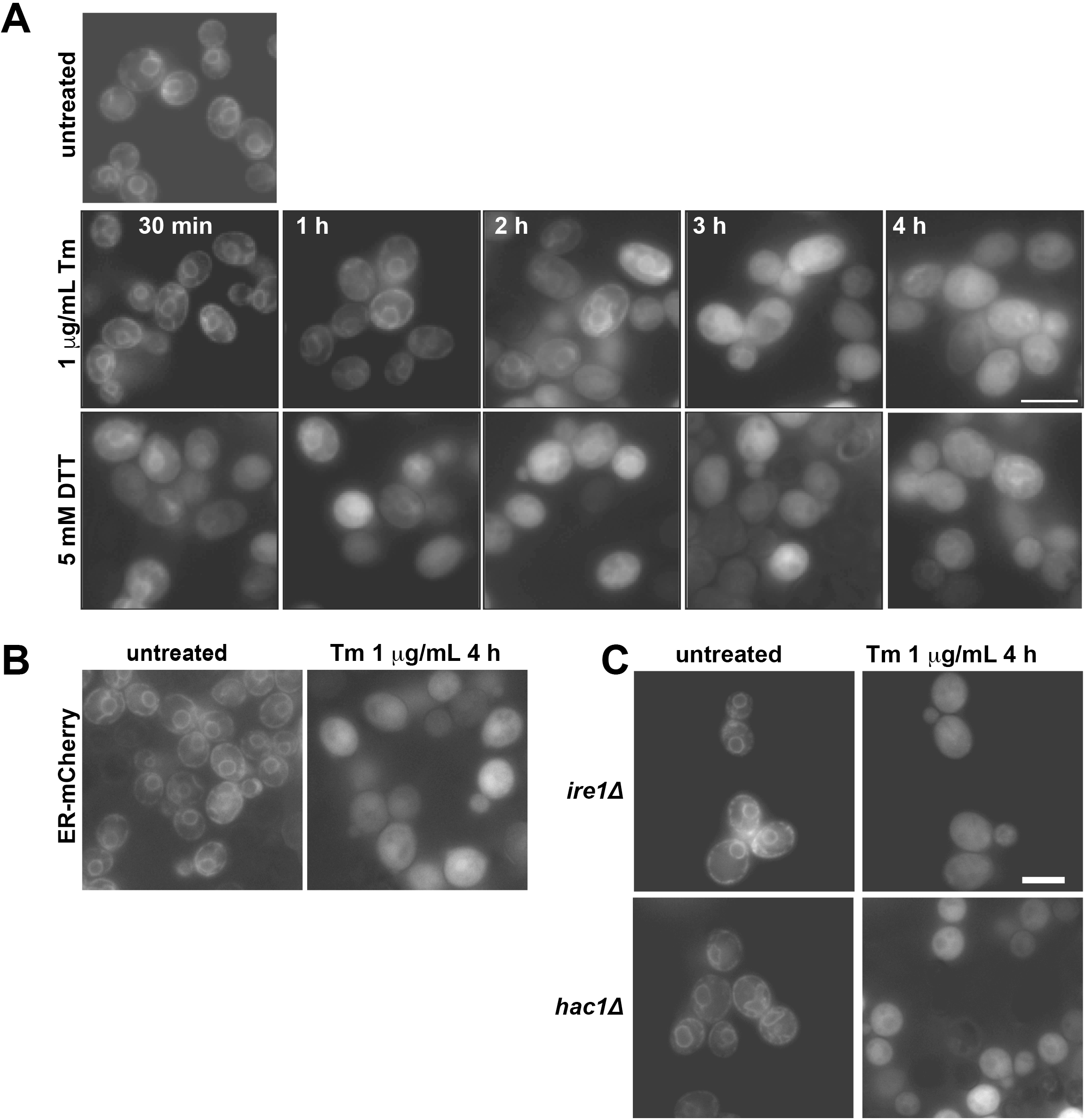
ER-sfGFP accumulates in the cytosol during acute misfolded secretory protein stress. (A) Treatment of strains expressing ER-sfGFP with secretory protein misfolding agents Tm or DTT lead to localization of ER-GFP to the cytosol. (B) Cytosolic localization also occurs for an ER-mCherry with Tm treatment. (C) The key UPR effectors are not linked to cytosolic accumulation of ER-sfGFP. *ire1Δ* and *hac1Δ* cells still exhibit significant cytosolic ER-sfGFP with Tm treatment. Scale bars = 5 μm.

As a working hypothesis, we speculated that ER stress was impairing efficiency of translocation of nascent proteins into the ER. Different signal sequences (SSs) translocate with distinct efficiencies during homeostasis and stress (8, 9). Therefore, we tested whether the choice of SS affected localization of ER-sfGFP during stress. We tried six different SSs (see Table 1), including SSs of proteins that translocate co-translationally (Dap2), post-translationally (CPY, Pdi1) or both (Kar2) (18). The mechanism of Scj1 translocation is unknown. Two different lengths of mature domains of the proteins immediately following the SS were also included (+3 or +10) because Levine et al. reported that including a part of the mature domain of the protein increased translocation efficiency of reporters during unstressed conditions (8). We found that some SSs, especially Pdi1 and CPY, localized ER-sfGFP less well to the ER, even under unstressed conditions. However, all of the constructs localized to the cytosol after 2 h of Tm stress (Fig. 2 A).

**Table 1.**
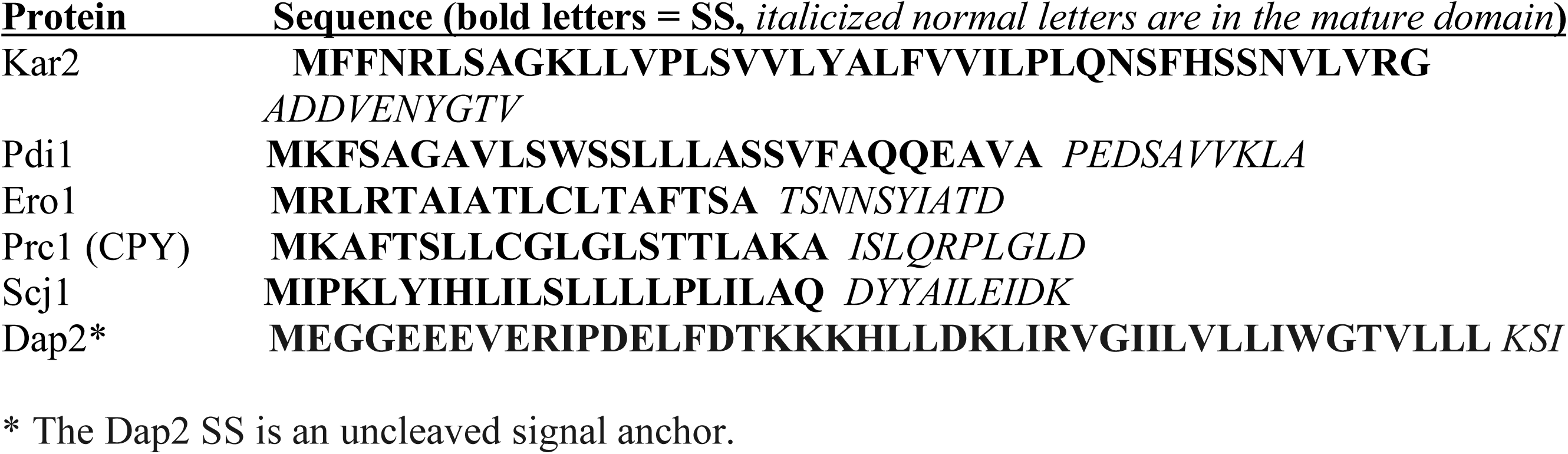
*S. cerevisiae* Protein Signal Sequences Used in this Study

**Figure 2.**
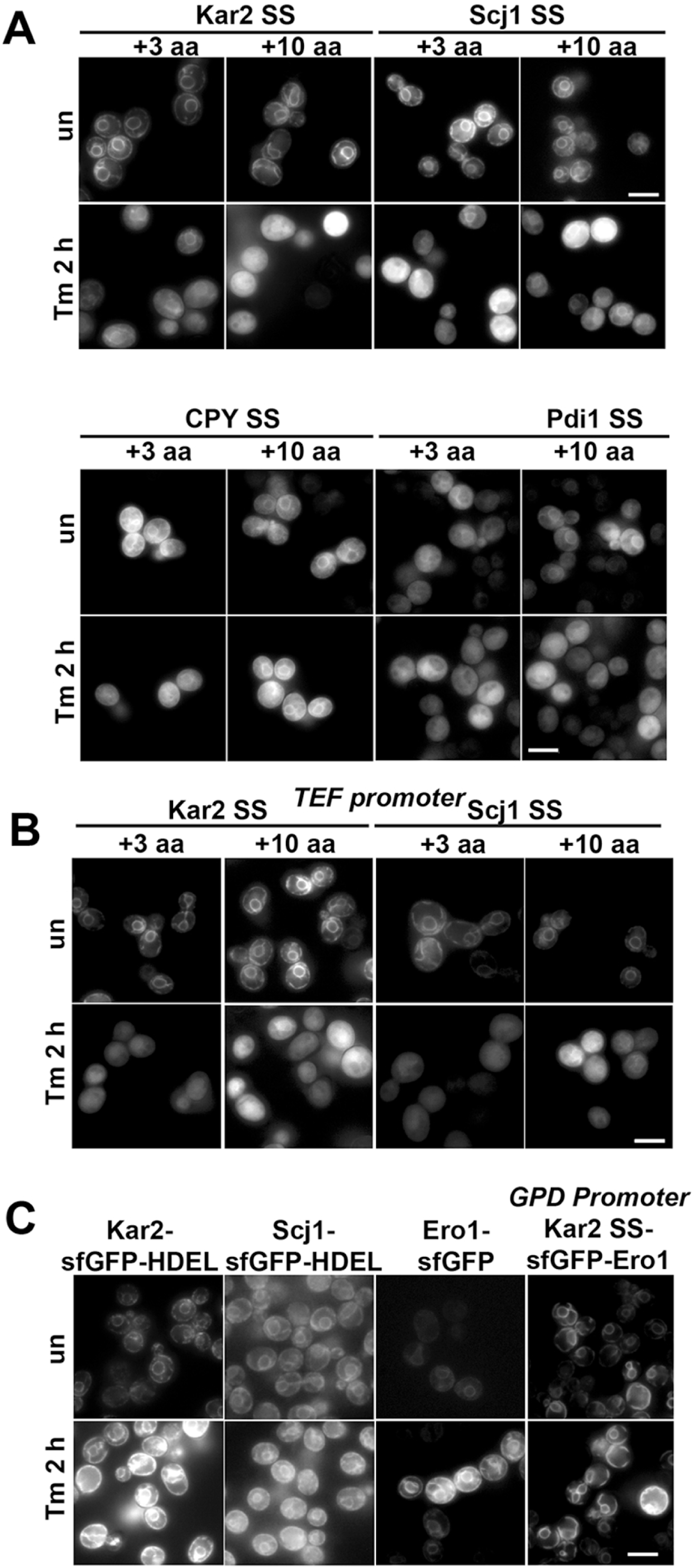
Different SSs and expression levels fail to maintain ER localization of ER-sfGFP reporters. (A) ER-sfGFP reporters with a variety of different secretory protein SSs accumulate in the cytosol with Tm treatment (5 μg/mL). Addition of 3 or 10 amino acids (+3 or +10 a.a.) of the mature domain after the SS do not improve ER localization with stress. In some cases, cytosolic localization occurs even in unstressed cells (CPY SS and Pdi1 SS). (B) Decreased expression levels with a TEF promoter do not prevent cytosolic localization during Tm stress of ER-sfGFP reporters with either the Kar2 or Scj1 SS in low copy plasmids. (C) Resident ER chaperones fused to sfGFP correctly localize to the ER during acute misfolded protein stress. sfGFP can be placed between the SS and the mature protein domain and Ero1-sfGFP ER localization remains robust during Tm stress. Scale bars = 5 μm.

Another consideration was that the ER-sfGFP constructs were expressed from multicopy plasmids with a constitutive robust promoter. Perhaps high ER-sfGFP expression overwhelms the translocation machinery during acute ER stress? To test this hypothesis, we decreased expression of ER-sfGFP using a weaker *TEF* promoter (19, 20). However, Tm still induced cytosolic accumulation of the ER reporter (Fig. 2 B).

We have previously tagged endogenous full length ER-chaperones with sfGFP (16) and did not notice significant cytosolic accumulation during acute ER stress. We decided to test other full length protein fusions for localization. Additional fusions to endogenous genes were made by homologous recombination. In unstressed cells, ER localization was robust (Fig. 2C). Following Tm treatment, we observed a substantial increase in fluorescence intensity, consistent with upregulation of the UPR targets Kar2, Scj1, and Ero1 (21). However, no significant cytosolic accumulation was apparent with stress (Fig. 2 C). We noted that the position of sfGFP in the fusions was at the end of relatively large proteins, raising the possibility that the translocation machinery might interact more robustly with the sequences of native ER proteins and enhance translocation efficiency during ER stress. Possibly, in these fusions, sfGFP might not engage the translocation machinery until after hundreds of amino acids already had been translocated and folded. In contrast, the sfGFP in the ER-sfGFP reporter would engage the translocation machinery immediately after the SS. As GFP was originally a cytosolic protein from jellyfish (22), it is possible that GFP lacks key protein sequences required for ER entry during stress. To test this hypothesis, we engineered sfGFP between the Kar2 SS and the mature domain of the robustly ER-localized resident ER protein Ero1. During Tm treatment, ER localization of the engineered construct was at least as robust as with the COOH-tagged endogenous Ero1-sfGFP (Fig. 2C). Therefore, the presence of sfGFP on a resident ER protein or immediately after the SS did not, in itself impact cytosolic localization during stress. Taken together, two different ER-targeted FPs fail to correctly localize to the ER during acute stress, while tagged resident ER proteins appear to be unaffected in their localization during ER stress.

Next, we asked whether the cytosolic pool of ER-sfGFP resulted from a translocation defect, as had been previously suggested. During translocation, many ER SSs are cleaved and this includes the Kar2 SS (23). Therefore, we investigated whether the Kar2 SS of ER-sfGFP was cleaved or uncleaved in stressed cells. Using a 12% tricine gel with sufficient resolution, we compared the sizes of two ER-sfGFP reporters relative to an unprocessed cytosolic sfGFP-HDEL in unstressed and stressed cells. One ER-sfGFP contained the cleavable Kar2 SS replaced with the ER-targeting uncleaved Dap2 signal anchor (48 aa or ∼5 kDa) that is similar in size to the uncleaved Kar2 SS+3 (45 aa) (Fig. 3 A). In stressed and unstressed conditions, ER-sfGFP migrates much faster than the Dap2 SS ER-sfGFP confirming that the Kar2 SS must be cleaved, consistent with trafficking to the translocon and cleavage by signal peptidase. In contrast, ER-sfGFP migrates slightly slower than the unmodified cytosolic sfGFP, which should be 8 amino acids (aa) shorter (∼1 kDa). The results of the immunoblot argue that ER-sfGFP must be translocated to be cleaved during ER stress before then accumulating in the cytosol.

**Figure 3.**
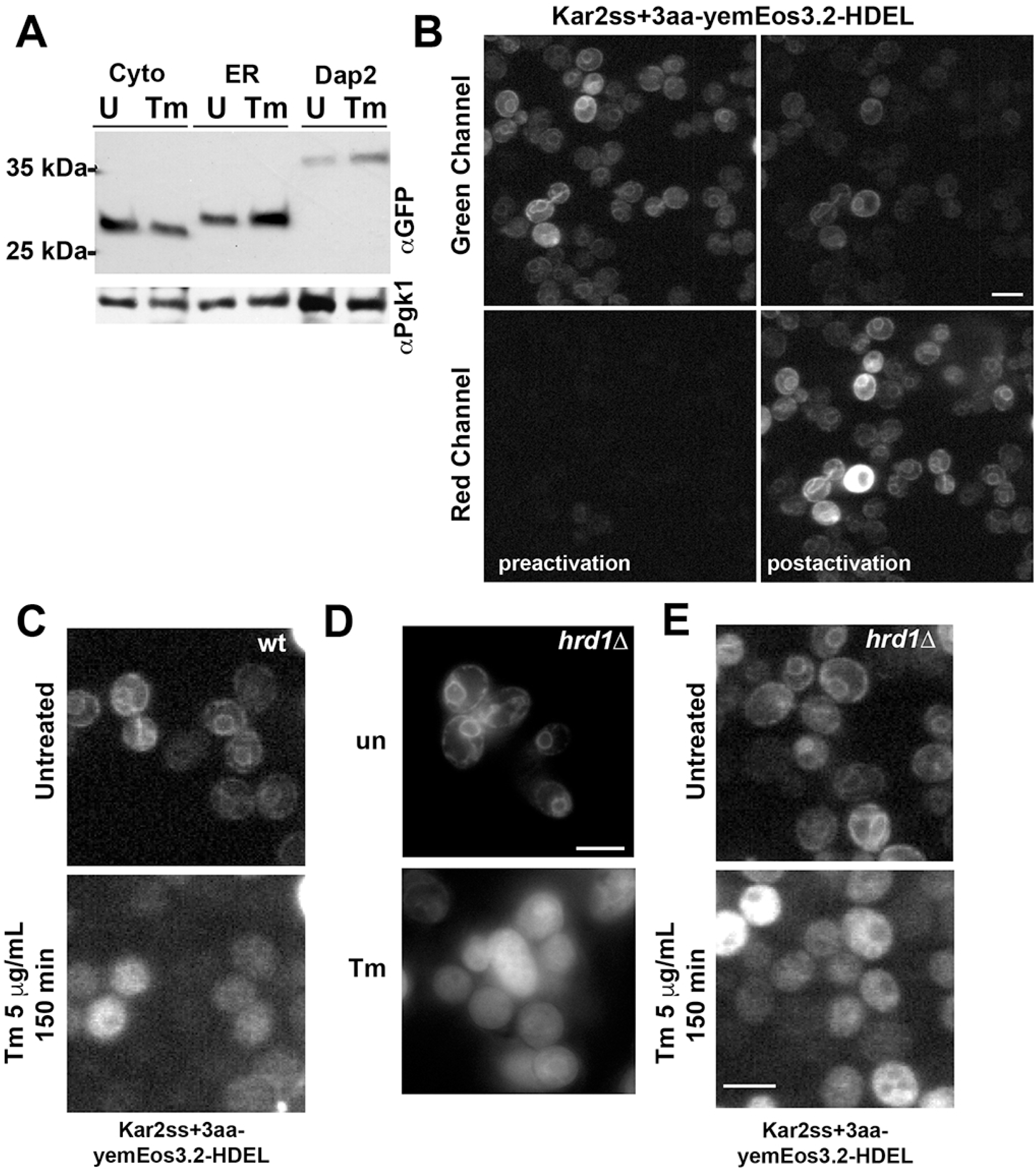
ER reporter that accumulates in the cytosol originated in the ER, but does not appear to rely on ERAD for relocalization. (A) Tm treatment does not significantly alter processing of secretory proteins. No size shifts are apparent for ER-sfGFP relative to the slightly smaller and unprocessed cytosolic GFP or the signal anchored Dap2, suggesting SS cleavage (ER) or lack of cleavage (the uncleaved Dap2 signal anchor) are not significantly impacted during acute stress. Pgk1 serves as a loading control. (B) A yeast codon optimized mEos3.2 (yemEos3.2) replaces sfGFP in the ER reporter and can be permanently converted from green to red when cells are briefly photoactivated with a 405 nm laser. (C) Reporters in cells were photoactivated to red and then stressed with Tm. Cytosolic accumulation of the ER reporters occur in both wt and ERAD-defective cells (*hrd1Δ*). In contrast, inserting a 24 a.a. GS linker into the ER reporter maintains ER localization of the photoconverted red population during Tm stress. (D) Cytosolic localization appears to be independent of ERAD-L. Cells inhibited from ERAD of luminal clients (*hrd1Δ*) still exhibit cytosolic localization of ER-sfGFP during Tm treatment. (E) Similar results are observed for optically highlighted (photoconverted) ER-yemEos3.2 in *hrd1*Δ cells, which still relocates from the ER lumen to the cytosol following Tm treatment. Scale bars = 5 μm.

This result raised the possibility that ER-sfGFP might completely translocate into the ER and then be retrotranslocated back into the cytosol, potentially by the ER associated degradation (ERAD) pathway (3, 5). We tested this hypothesis using two approaches. First, we exploited the power of photoactivatable proteins that can be permanently optically highlighted by laser light converting the protein from a green to a spectrally distinct red species. Using yeast codon optimized mEos3.2 (yemEos3.2) (24), we demonstrated that the ER-targeted FP can be robustly converted from green to red in the yeast ER (Fig. 3 B). Surprisingly, Tm treatment of cells with photoconverted ER-mEos3.2 revealed that initially ER-localized reporter *was relocalizing to the cytosol during stress* (Fig. 3 C). Thus, some form of ERAD could potentially move the ER reporters from the ER lumen to the cytosol. This interpretation was problematic.

The first concern was whether the reporter was incorrectly folded. The standard model of ERAD-L, the pathway by which luminal proteins are recognized and retrotranslocated to the cytoplasm by the ERAD machinery, holds that the luminal target proteins are misfolded and recognized by ER chaperones, such as Kar2 or BiP (in metazoans) or BiP-associated J protein co-factors (25). Given that the FP reporters were fluorescent in the ER argues that the FPs must have first folded into a beta barrel, which is essential to fluorophore formation (26). Therefore, to be an ERAD target, the ER FP reporters must be somehow recognized as either unfolded or might somehow be unfolded directly by chaperones. This latter model has some precedence for some folded viral proteins and bacterial toxins that ER chaperones have been observed to engage and apparently unfold (27). We will return to this question in Figure 4.

**Figure 4.**
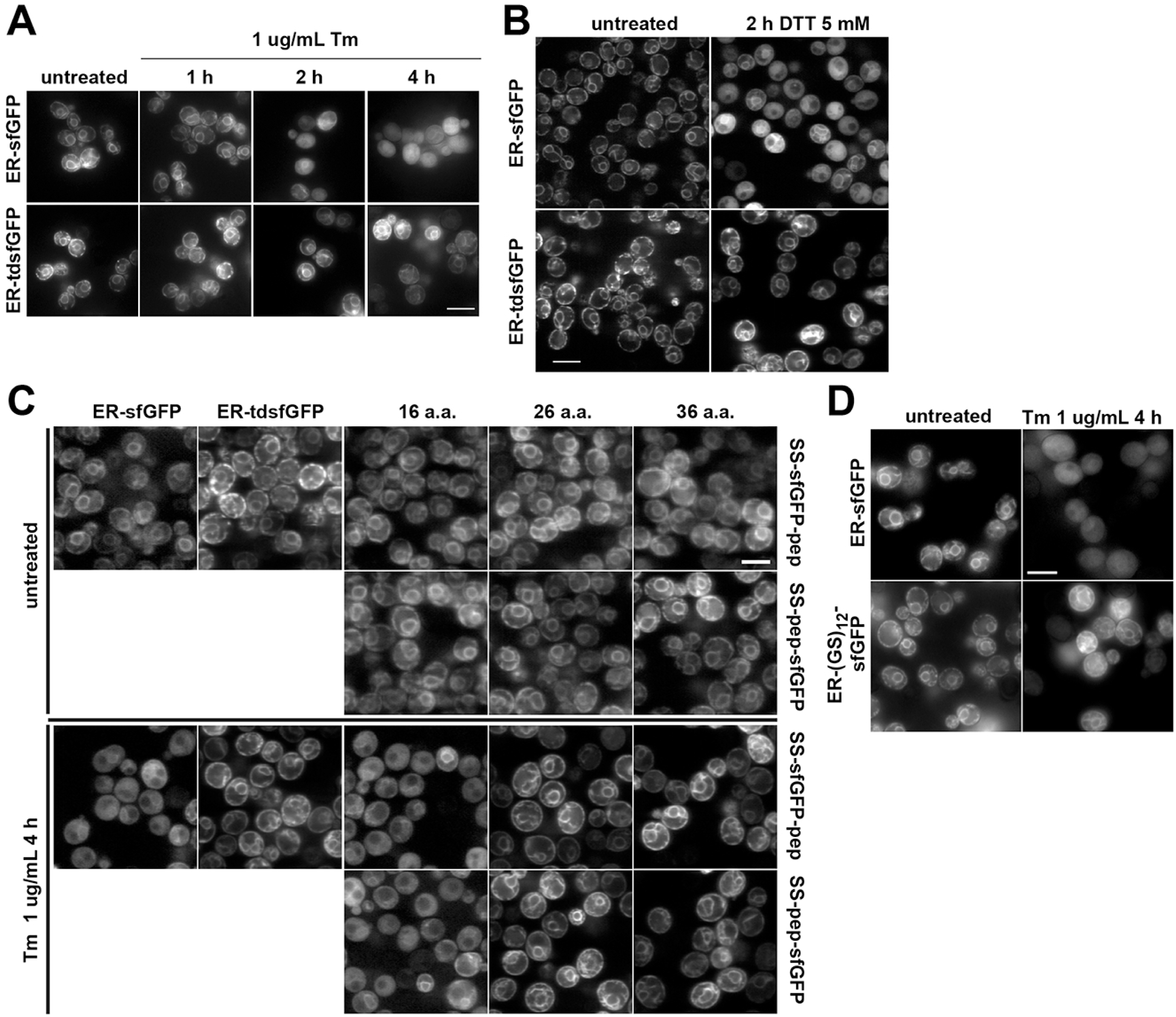
ER localization during misfolded secretory protein stress is improved with increased reporter size. (A) Yeast expressing ER-sfGFP or ER-tdsfGFP were treated with 1 μg/mL Tm for indicated times and imaged by fluorescence microscopy. (B) As in a, but treated with 5 mM DTT. Unlike the other fluorescence micrographs in Fig. 4, part b was imaged using confocal microscopy. (C) Yeast expressing the indicated constructs were either untreated or treated with Tm for 4 h. Additions of 26 and 36 a.a. of sfGFP, as a linker, resulted in robust ER localization during Tm treatment. The slightly shorter 16 a.a. linker did not. (D) A 24 a.a. non-GFP linker (Gly-Ser)_12_ also robustly maintained ER localization of a SS-pep-sfGFP reporter during 1 μg/mL Tm after 4 h. Scale bar = 5 μm.

The second concern is that most ERAD-L clients are destroyed by the proteasome and do not refold in the cytosol (5). The retrotranslocation of luminal proteins into the cytoplasm is coupled to ubiquitination and subsequent targeting to the proteasome for degradation (28). Bacterial toxins retrotranslocated by ERAD-L, the form of ERAD that recognizes and retrotranslocates soluble secretory proteins, appear to escape destruction in part by a low lysine content, which decreases opportunities for ubiquitination (27). FPs contain multiple lysines (20 out of 239 residues in sfGFP, for example) and are susceptible to proteasome degradation when FPs are fused to misfolded proteins (29). Taken together, either FPs are retrotranslocated to the cytoplasm by an ERAD mechanism similar to those utilized by viruses and bacterial toxins or ERAD is not the mechanism.

The ERAD-L pathway requires the membrane protein Hrd1, which is hypothesized to form a channel from the ER lumen to the cytosol (28). Deletion of *hrd1* disrupts ERAD-L (30, 31), but did not prevent relocalization of ER-sfGFP (Fig. 3 D) or photoactivated ER-yemEos3.2 (Fig. 3 E) during Tm treatment. Together, these experiments argue that ER-sfGFP enters the ER, undergoes SS cleavage and correct folding, and then is retrotranslocated by a mechanism that appears to be distinct from ERAD-L. Therefore, we have termed this phenomenon “ER reflux.”

### A link between ER reflux and Heat Shock

An important question is whether ER reflux can be stimulated by other acute misfolded protein stresses. We examined whether heat shock can trigger ER reflux and observed no obvious cytosolic accumulation of ER-sfGFP even after 4 h of incubating cells at a stressful 40°C (Fig. S1 A). We also tested whether heat shock might exacerbate or accelerate ER reflux and found the opposite. Instead, cells treated with Tm and grown at 40°C exhibited protection against Tm-induced ER reflux (Fig. S1 A)(32). The heat shock response up regulates expression of 165 genes and it is unclear which specific proteins are implicated in ER reflux (33).

### Engineering an ER reporter resistant to ER reflux

To enable study the ER environment during acute ER stress, we sought to determine how ER reflux might be circumvented to maintain localization of reporters inside the ER. We considered the results in Figure 2 with the various reporters and the ability of tagged endogenous ER proteins to remain in the ER during acute stress. We noted that fusions to full-length reporters remained ER localized, which suggested the ER proteins might contain retention information or that retention might be due to size. All of the fusions were significantly larger molecules than GFP alone, ∼5 nm (34). We chose to test the latter hypothesis and doubled the size of our reporter by making a tandem dimer (td) fusion. Consistent with the size-dependent hypothesis, the new ER-tdsfGFP exhibited robust ER localization during acute ER stress with Tm or DTT (Fig. 4 A and B). The larger tdsfGFP also improved ER localization in unstressed cells when the Kar2 SS was replaced with the less robustly ER-localized Pdi1 SS (Fig. S2 A and B).

We were curious whether the bulky size of the complete additional GFP β-barrel was necessary to block ER reflux or whether a shorter truncated peptide would be sufficient to confer resistance to ER reflux. Treatment with Tm led to significant cytosolic accumulation of Kar2 SS ER-sfGFP fused to the first 16 aa of sfGFP at either the COOH terminus of sfGFP or in between the cleaved SS and fused to the NH2 start of the mature domain of Kar2 (Fig. 4 C). Surprisingly, we found that addition of a peptide of 26 aa or more of the sfGFP sequence was sufficient to prevent ER reflux and maintained ER localization regardless of whether the peptide was placed before the mature domain of Kar2 or at the COOH-terminus of the full length sfGFP (Fig. 4 C). ER localization was comparable to ER-tdsfGFP. Given the lack of a positional requirement and the fact that jellyfish GFP is a cytosolic protein, we hypothesized that only the size of the peptide, not the sequence was important for protection. To test this, we inserted a 24mer consisting of GS_12_ between the SS and the mature sfGFP. This short peptide proved sufficient to protect against ER reflux of ER-sfGFP (Fig. 4 D). Finally, we confirmed that pre-existing protein was retained in the ER by adding the SS+24 aa linker to yemEos3.2 and stressing cells labeled with photoconverted protein (Fig. S2 C). Thus, a short peptide in *cis* can maintain ER-sfGFP localization during acute misfolded protein stress. This should prove extremely helpful for monitoring ER morphology and retaining biosensors in the ER during acute misfolded protein stress conditions.

One important parameter for assays of changes in the organization of the ER lumen is viscosity or crowdedness. This parameter can be measured with Fluorescence Recovery after Photobleaching (FRAP) and an inert reporter, such as sfGFP (35, 36). The need to increase the size of ER-GFP to maintain ER localization made it unclear if reporter mobility would be significantly altered by the increased size. For, ER-tdsfGFP, we predicted that linking two sfGFP molecules together (which should result in a relatively large ∼10 nm protein) would likely exhibit low mobility. We compared the mobilities of ER-tdsfGFP and ER-sfGFP-(GS)_12_ reporters in environmental viscosity FRAP assays. In a cell, ER-sfGFP diffuses at 2.3 µm^2^/s (Figure 5). The Stokes-Einstein equation predicts that doubling the size of ER-sfGFP should decrease the diffusion coefficient, *D*, by one half (37). In contrast, ER-tdsfGFP exhibited an extremely low *D* value of 0.5 µm^2^/s or one fifth of ER-sfGFP. This low value could arise due to an unanticipated interaction with a resident ER protein or more likely due to the relative narrowness of the yeast ER lumen (∼10-76 nm with a mean diameter of 37.9 nm)(38). Regardless, the low starting *D* value results in a narrow dynamic range for using the reporter to detect changes in ER viscosity. In contrast, the much shorter construct, SS-(GS)_12_-sfGFP-pep, exhibited indistinguishable diffusion properties relative to ER-sfGFP (Figure 6B). Thus, addition of a short peptide is sufficient to maintain localization and functionality of FP reporters and biosensors in the yeast ER.

**Figure 5.**
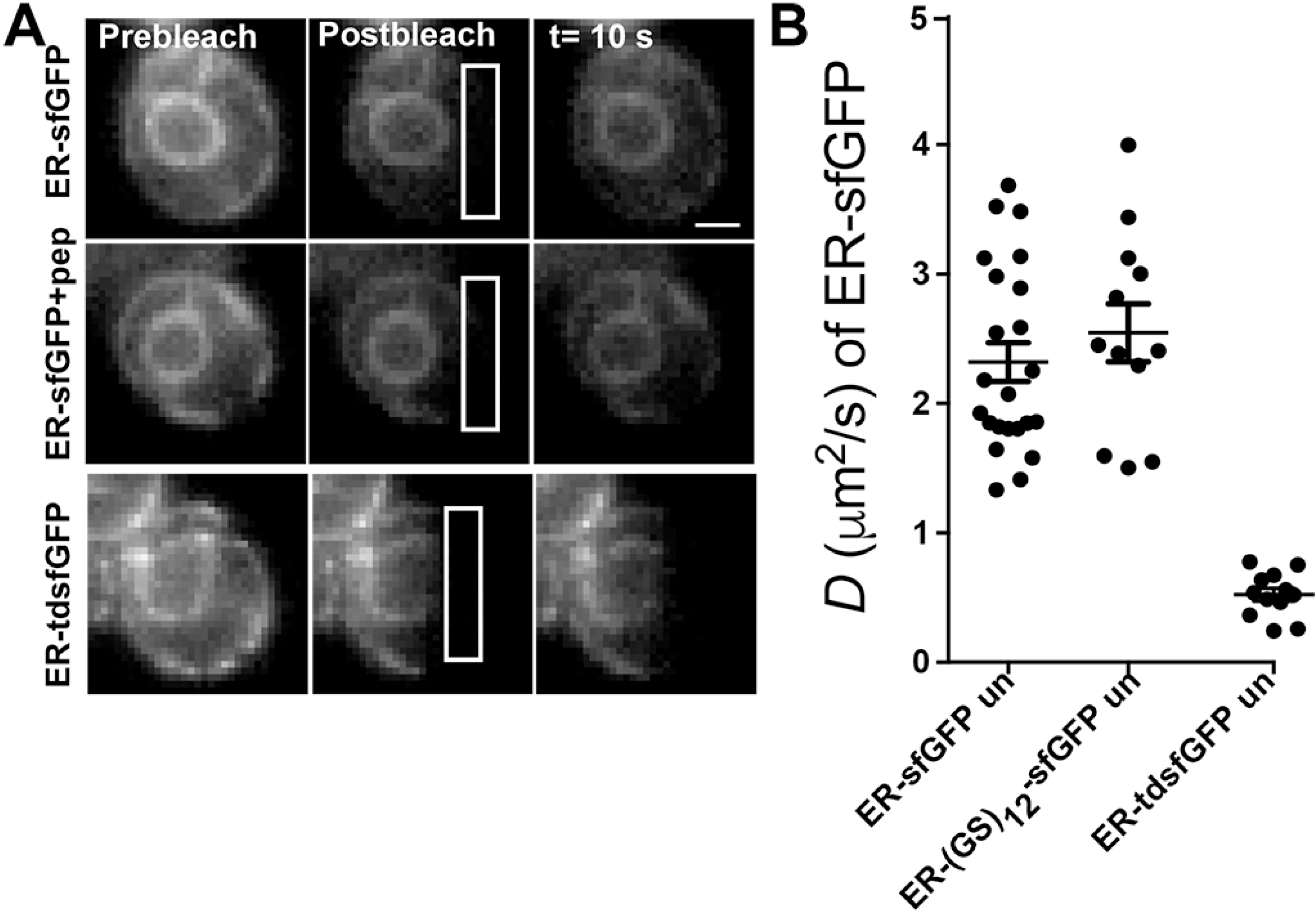
ER-sfGFP-pep is a neutral environmental reporter of the ER lumen. (A) FRAP time series of yeast expressing ER-sfGFP-pep (the +26 aa construct from Fig. 5) or ER-tdsfGFP. White ROIs indicate photobleach regions. (B) Plot of *D* values for different ER-sfGFP constructs. Each closed circle marks the *D* value of a single cell. Bars within the dots indicate mean *D* value and S.E.M. n ≤. Scale bar = 1 μm.

## DISCUSSION

In this study, we describe a novel phenomenon in which a small soluble ER-localized protein is expelled from the ER lumen during the acute stressful accumulation of misfolded secretory proteins. Our photoactivation experiments convincingly distinguish this process from some form of failed translocation of nascent secretory proteins. The *hrd1Δ* mutant results strongly suggest ER reflux is distinct from standard models of ERAD. Equally importantly, ER reflux moves a correctly folded protein from the ER to the cytosol and where the protein is also functionally folded. We can draw this conclusion because FPs must form the elaborate β-barrel structure to form the fluorophore and for fluorescence to occur (39). The result with mEos3.2 follows a folded FP in the ER lumen that is converted and then the converted fluorescent form appears in the cytosol. Currently, we cannot rule out the possibility that the protein unfolds to move from the ER lumen to the cytosol and then refolds. However, we consider this possibility unlikely because of our findings on the effect of size or protein primary sequence length. It is not difficult to envision a model in which ER reflux machinery could sort proteins by globular size (e.g. hydrodynamic radius), it’s less obvious why or how the cell would distinguish unfolded proteins that differ in length by 10 amino acids.

ER reflux is triggered by acute pharmacologic stresses that induce global misfolding of secretory proteins, but not a cytosolic stress, such as heat shock. Yet, neither an ERAD-L deletion mutant, *hrd1Δ* (Fig. 3 D), nor deletion of the key UPR pathway components Ire1p or Hac1p appear to cause or prevent ER reflux (Fig. 1 C), respectively. Together, these observations suggest that yeast possess a novel mechanism for detecting and responding to acute accumulation of misfolded secretory proteins in the ER. However, the release of correctly folded proteins below a molecular size cutoff suggests that ER reflux may not necessarily be a mechanism for clearing the ER lumen of misfolded proteins to restore homeostasis. Most relatively small ER proteins in yeast are integral membrane proteins, which do not appear to be subject to the ER reflux pathway (Feroz Papa and Aeid Igbaria, personal communication).

The role of ER reflux remains unclear. Reflux could function to remove otherwise correctly folded and functional small secretory proteins and consequently block fundamental yeast processes, including mating/conjugation (i.e. Mf(alpha)2 and Mf(alpha)1)(also see Table 2). A curious possibility that we cannot currently rule out is that FPs in yeast could become targets of ER chaperones during conditions of misfolded secretory protein stress. We do not observe this in mammalian cells by FRAP analysis (29, 40, 41). We have not tested this possibility by co-IP in actively stressed cells as the previous experiments we performed were in homeostatic cells. The FRAP experiments are further confounded by the impact of chaperone binding of FPs-namely that the FPs would likely unfold, become dark, and be undetectable by fluorescence microscopy. One potential mechanism to drive interactions between FPs and chaperones, such as Kar2p/BiP, is the dramatic UPR induced upregulation of Kar2p levels. At steady state, a significant fraction of Kar2p/BiP is occupied (16, 41, 42). Increased levels of BiP/Kar2p could increase the probability of interactions with FPs that might have low affinity chaperone binding sites. However, this model seems unlikely, because we still observe ER reflux in UPR impaired mutants (Fig. 1C). Another challenge with the FP misfolding model is that both a tandem dimer and an FP fused with either a short FP-derived peptide or a repeating GS peptide of a minimal length are sufficient to prevent ER reflux. It’s unclear how a short flexible peptide would protect FP fusions from chaperone binding, unless the peptide somehow sterically blocks a chaperone binding site.

**Table 2.** Candidate Soluble Yeast Proteins Potentially Subject to ER reflux

| **<u>Systematic Name</u>** | **<u>Gene Name</u>** | **<u>Protein Length (amino acids)</u>** |
| --- | --- | --- |
| YGL089C | MF(Alpha)2 | 120 |
| YPL187W | MF(Alpha)1 | 165 |
| YGL258W | VEL1 | 206 |
| YDR304C | CPR5 | 225 |
| YOL088C | MPD2 | 277 |

As a consequence of these studies, it is clear that biosensors and FP reporters require validation of functionality and expected localization in cells under the conditions to be explored. This represents another parameter, in addition to the often overlooked impact of different cell environments on biosensor folding and function (43-45). In this study, the size of some proteins may render the proteins susceptible to relocalization, during misfolded secretory protein stress, for example. Less obvious is the corollary of our findings, small resident ER proteins tagged with GFP or even a 10 amino acid tag could dramatically alter the ability of proteins to correctly relocalize in response to environmental stimuli. For example, Shao and Hegde described an ER-targeting and translocation pathway for small (<160 a.a.) secretory proteins (46). Small secretory proteins traffic post-translationally from the cytosol to the Sec61 translocon via an interaction with calmodulin. Addition of a GFP or epitope tag would lengthen these proteins and switch these proteins over to the co-translational Signal Recognition Particle-dependent targeting pathway.

Ultimately, our data argue for caution when using GFP, other FPs or epitope tags to localize or characterize proteins. Raising antibodies against uncharacterized proteins remains an important component of basic cell biology research. Using tagged molecules, an undeniably powerful tool, to characterize proteins requires extreme caution and careful validation that tags do not perturb protein behavior and accurately reflect protein behavior in cells.

## MATERIALS AND METHODS

### Drugs

Stock solutions of DTT (1 M in water; Fisher Scientific, Pittsburgh, PA) and Tm (5 mg/ml in DMSO; Calbiochem, La Jolla, CA) were prepared and used at the indicated concentrations and times indicated.

### Strains and Cell Growth

See Supplementary Table 1 for all yeast strains used in this study. All strains were derived from By4741 (*MAT*α *his3Δ0 leu2Δ0 met15Δ0 ura3Δ0*) with transformed plasmids selected by dominant drug markers. The *hrd1Δ* strain was obtained from Dr. Ian Willis (Albert Einstein College of Medicine, New York). Kar2-sfGFP-HDEL strain was made previously (16). All yeast strains were grown in synthetic complete media supplemented with appropriate amino acids. Yeast strains were grown at 30°C. Yeast strains were grown overnight to early log phase (OD600 nm ≈ 0.5) for analysis.

### Plasmid Constructions

Please see Supplemental Information for plasmid construction information.

### Heat Shock Assay

Yeast cells were grown in the synthetic complete media supplemented with appropriate amino acids at 25°C to the early log phase. Cells were either treated with Tm or untreated, and cultured at 30°C or 40°C, followed by fluorescence acquisition with time.

### Fluorescence Microscopy

After incubation with indicated stresses or stressors, log-phase cells were placed in 8-well Lab-Tek chambers (ThermoFisher Scientific, Waltham, MA) and allowed to settle for 5 min before imaging. Cells were imaged in SC complete with appropriate selection components on an Axiovert 200 wide-field fluorescence microscope (Carl Zeiss MicroImaging, Inc., Thornwood, NY) with a 63X/1.4 NA oil immersion objective lens, a Retiga-2000 camera (QImaging, Surrey, BC Canada), and 470/40 nm excitation, 525/50 nm emission bandpass filter for GFP, 565/30 nm excitation, or 565/30 nm excitation, 620/60 nm emission bandpass filter for mCherry. Images were acquired with QCapture software. Confocal images were acquired with a Zeiss LSM-5 LIVE microscope with Duoscan attachment (Carl Zeiss MicroImaging) with a 63X, N.A. 1.4 oil objective and a 489 nm, 100 mW diode laser with a 500-550 nm bandpass filter for GFP or a 561 nm diode laser with a 565 nm longpass filter for mCherry. Assessment of FP localization was determined visually. Due to the small size of yeast, scatter, and autofluorescence, cells were scored simply for a nuclear and peripheral ER pattern versus substantial cytosolic and nuclear accumulation, as well as the frequent visibility of the yeast vacuole.

Photobleaching was also performed on the LSM-5 LIVE. FRAP experiments were performed by photobleaching a region of interest at full laser power of the 489-nm line and monitoring fluorescence loss or recovery over time. No photobleaching of the adjacent cells during the processes was observed. *D* measurements were made using an inhomogeneous diffusion simulation, as described previously (47, 48). Image analysis was performed with ImageJ (National Institutes of Health, Bethesda, MD) and composite figures were prepared using Photoshop CC2018 and Illustrator CC2018 software (Adobe Systems, San Jose, CA).

### Immunoblots

Early-log-phase yeast strains, untreated or treated with Tm, were pelleted and total protein extracted by alkaline lysis (49). Lysates were separated on 7.5% or 12% SDS–PAGE tricine gels, transferred to nitrocellulose membranes, and detected with anti-GFP (from Ramanujan Hegde, MRC Laboratory of Molecular Biology, Cambridge, United Kingdom), and horseradish peroxidase–labeled anti-rabbit (Jackson ImmunoResearch Laboratories, West Grove, PA).

### Statistical Analysis

Prism software (GraphPad Software, San Jose, CA) was used to compare the different conditions using two-tailed Student’s t-tests. For higher stringency, differences were not considered significant for *p* values >0.01.

## Acknowledgements

We thank Ian Willis, Robyn Moir, and Feng Guo (Albert Einstein College of Medicine) for reagents, assistance with experiments, and helpful conversations. We thank Ramanujan Hegde (Laboratory of Molecular Biology, Cambridge, UK) for the anti-GFP antibody, and the Einstein Analytical Imaging Facility for use of the Zeiss Duoscan. We thank Feroz Papa (University of California, San Francisco) for sharing key data from his lab’s studies and for helpful discussions that stimulated this study.

This work is supported by grants from the National Institute of General Medical Sciences (NIGMS)(1R01GM10599-01)(E.L.S.).

The authors declare no competing financial interests.

## Abbreviations

aa: amino acids
bp: base pairs
*D*: diffusion coefficient
DTT: Dithiothreitol
ER: Endoplasmic Reticulum
ERAD: Endoplasmic Reticulum Associated Degradation
FP: Fluorescent Protein
FRAP: Fluorescence Recovery after Photobleaching
GFP: Green Fluorescent Protein
MFI: Mean Fluorescence Intensity
QC: Quality Control
sfGFP: Superfolder GFP
SS: Signal Sequence
td: tandem dimer
Tm: Tunicamycin
UPR: Unfolded Protein Response

## MATERIALS AND METHODS

### Plasmid Constructions

SfGFP-HDEL with the first 135 base pairs of the Kar2p coding sequence, i.e. Kar2p signal sequence (SS) + the first 3 amino acids of the mature domain (Kar2pSS+3aa) was described previously (16). SfGFP-HDEL with the coding sequence of the Kar2p signal sequence plus 10 additional amino acids (Kar2pSS+10aa) was made by inserting the first 156 base pairs of the Kar2p coding sequence into the *Bgl*II/*Age*I sites of sfGFP-HDEL-N1 plasmid (46, 47) using the following primers:

Forward, GATCAGATCTCTAAAAATGTTTTTCAACAGAC

Reverse, GATCACCGGTCCAACAGTTCCGTAGTTTTC

The Kar2p+10aa-sfGFP-HDEL and Kar2p+3aa-sfGFP-HDEL fragments were subsequently amplified using the following primers to introduce yeast Kozak sequences:

Forward, GATCACTAGTCTAAAAATGTTTTTCAACAGAC

Reverse, GATCGGATCCTTACAATTCATCGTG

The resulting fragments were digested and cloned into the *Spe*I/ *BamH*I site of pRS415-GPD (a generous gift from Elizabeth Miller, Columbia University, New York, NY).

sfGFP-HDEL with Kozak sequence and the Cpy1p signal sequence plus 3 or 10 additional amino acids (Cpy1pSS+3aa/Cpy1pSS+10aa), Pdi1p signal sequence plus 3, 10 additional amino acids (Pdi1pSS+3aa/Pdi1pSS+10aa), or Dap2p signal sequence plus 3 additional amino acids (Dap2pSS+3aa) was made by inserting corresponding coding sequences into the *Bgl*II/*Age*I sites of sfGFP-HDEL using the following primers:

Cpy1pSS+3aa

Forward, GATCAGATCTCTAAAAATGAAAGCATTCACC

Reverse, GATCACCGGTCCCAATGAGATGGCCTTAG

Cpy1pSS+10aa

Forward, GATCAGATCTCTAAAAATGAAAGCATTCACC

Reverse, GATCACCGGTCCATCTAGACCCAACGG

Pdi1pSS+3aa

Forward, GATCAGATCTCTAAAAATGAAGTTTTCTGCTG

Reverse, GATCACCGGTCCCTCTTGTTGGGCGAAAAC

Pdi1pSS+10aa

Forward, GATCAGATCTCTAAAAATGAAGTTTTCTGCTG

Reverse, GATCACCGGTCCGGAGTCTTCAGGGGCCAC

Dap2pSS+3aa

Forward, GATCAGATCTCTAAAAATGGAAGGTGGCGAAG

Reverse, GATCACCGGTCCGTGAGGTATACTTTTTAGCAAC

These ER targeted sfGFP-HDEL fragments were subsequently amplified using the following primers:

Forward, GCAAATGGGCGGTAGGCG

Reverse, GATCGGATCCTTACAATTCATCGTG

The resulting fragment was digested with *Bgl*II and *BamH*I and cloned into the *BamH*I site of pRS415-GPD.

The tandem dimer sfGFP (tdsfGFP) or tdTomato with KDEL retrieval motif was inserted at the C-terminus on the sfGFP-N1 plasmid (37, 47) was modified to contain the yeast HDEL motif using the following primers (16):

Forward, GGACGAGCTGTACAAGGATGAATTGTAAGCG

Reverse, CGCTTACAATTCATCGTGGTACAGCTCGTCC

Kar2pSS+3aa-td-sfGFP-HDEL, Pdi1p+3aa-tdsfGFP-HDEL or Pdi1p+3aa-tdTomato-HDEL were made by inserting Kar2pSS+3aa or Pdi1pSS+3aa into the *Bgl*II/*Age*I site of tdsfGFP-HDEL or tdTomato-HDEL. The Kar2pSS-tdsfGFP-HDEL fragment was amplified using the following primers:

Forward, GATCACTAGTCTAAAAATGTTTTTCAACAGAC

Reverse, GATCGGATCCTTACAATTCATCGTG

The resulting fragment was digested and cloned into the *Spe*I/*BamH*I site of pRS415-GPD.

The Pdi1pSS+3aa-tdsfGFP-HDEL or Pdi1pSS+3aa-tdTomato-HDEL fragments were amplified using the following primers:

Forward, GCAAATGGGCGGTAGGCG

Reverse, GATCGGATCCTTACAATTCATCGTG

The resulting fragment was digested with *Bgl*II and *BamH*I and cloned into the *BamH*I site of pRS415-GPD.

The mCherry with KDEL retrieval motif inserted at the C-terminus on the mCherry-N1 plasmid was modified to contain the yeast HDEL motif using the following primers:

Forward, GGACGAGCTGTACAAGGATGAATTGTAAGCG

Reverse, CGCTTACAATTCATCGTGGTACAGCTCGTCC

Kar2pSS+3aa fragment was inserted into the *Bgl*II/*Age*I site of mCherry-HDEL. The Kar2pSS-mCherry-HDEL fragment was amplified using the following primers:

Forward, GATCACTAGTCTAAAAATGTTTTTCAACAGAC

Reverse, GATCGGATCCTTACAATTCATCGTG

The resulting fragment was digested and cloned into the *Spe*I/*BamH*I site of pRS415-GPD. Kar2pSS+3aa-tdsfGFP-HDEL series truncations were made by amplifying fragments using the following primers:

Forward, GCAAATGGGCGGTAGGCG,

Kar2pSS+3aa-sfGFP-16aa-HDEL

Reverse, GATCGCGGCCGCTTACAATTCATCGTGCAGGATGGGCACCACC

Kar2pSS+3aa-sfGFP-26aa-HDEL

Reverse, GATCGCGGCCGCTTACAATTCATCGTGGTGGCCGTTTACGTCG

Kar2pSS+3aa-sfGFP-36aa-HDEL

Reverse, GATCGCGGCCGCTTACAATTCATCGTG GCCCTCGCCCTCGCCG

Kar2pSS+3aa-sfGFP-46aa-HDEL

Reverse, GATCGCGGCCGCTTACAATTCATCGTGCTTCAGGGTCAGCTTG

Kar2pSS+3aa-sfGFP-157aa-HDEL

Reverse, GATCGCGGCCGCTTACAATTCATCGTGCTTGTCGGCGGTG

The fragments were inserted into the *Bgl*II/*Not*I sites of sfGFP-N1plasmid and were amplified using the following primers:

Forward, GATCACTAGTCTAAAAATGTTTTTCAACAGAC

Reverse, GATCGGATCCTTACAATTCATCGTG

The resulting fragments were digested and cloned into the *Spe*I/*BamH*I site of pRS415-GPD. Kar2pSS+3aa-sfGFP-HDEL series insertions were made by amplifying fragments using the following primers:

Forward, GATCACCGGTCGTGAGCAAGGGCGAG,

Kar2pSS+3aa-16aa -sfGFP-HDEL

Reverse, GATCACCGGTCCCAGGATGGGCACCACC

Kar2pSS+3aa-26aa -sfGFP-HDEL

Reverse, GATCACCGGTCCGTGGCCGTTTACGTCG

Kar2pSS+3aa-36aa -sfGFP-HDEL

Reverse, GATCACCGGTCCGCCCTCGCCCTCGCCG

The fragments were inserted into the *Age*I site of Kar2pSS-sfGFP-HDEL-N1plasmid and were amplified using the following primers:

Forward, GATCACTAGTCTAAAAATGTTTTTCAACAGAC

Reverse, GATCGGATCCTTACAATTCATCGTG

The resulting fragments were digested and cloned into the *Spe*I/*BamH*I site of pRS415-GPD. Yeast codon optimized sfGFP (yesfGFP) was purchased from GenScript (Piscataway, NJ). Kar2pSS-yeSfGFP-HDEL was made similarly as Kar2pss-sfGFP-HDEL.

**Figure S1.**
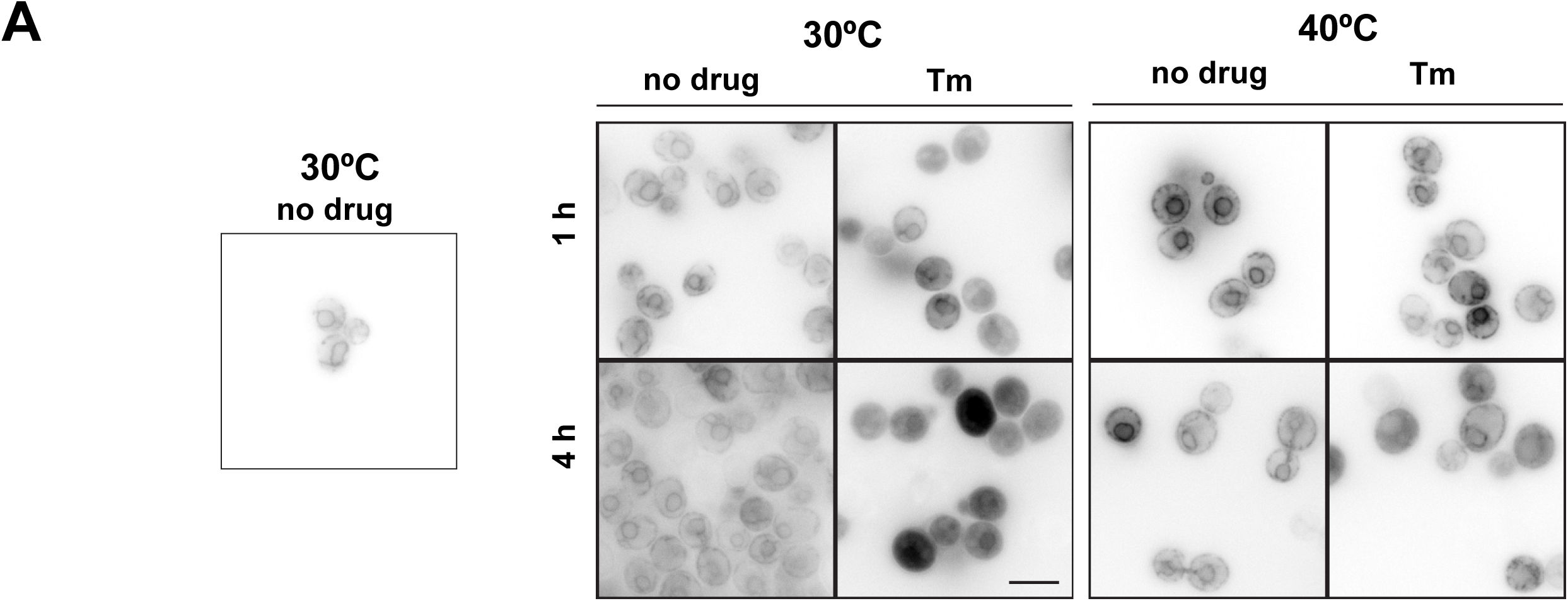
Heat Shock blocks ER reflux. (a) Cells were grown to early log phase at 30°C, diluted and then grown with or without 1 μg/mL Tm at the indicated temperatures, imaged with a widefield microscope, and then images were inverted for ease of visualizing the nuclear envelope and peripheral ER. Growth at 40°C in Tm protected against significant accumulation of ER-GFP in the cytosol. Scale bar = 5 μm.

**Figure S2.**
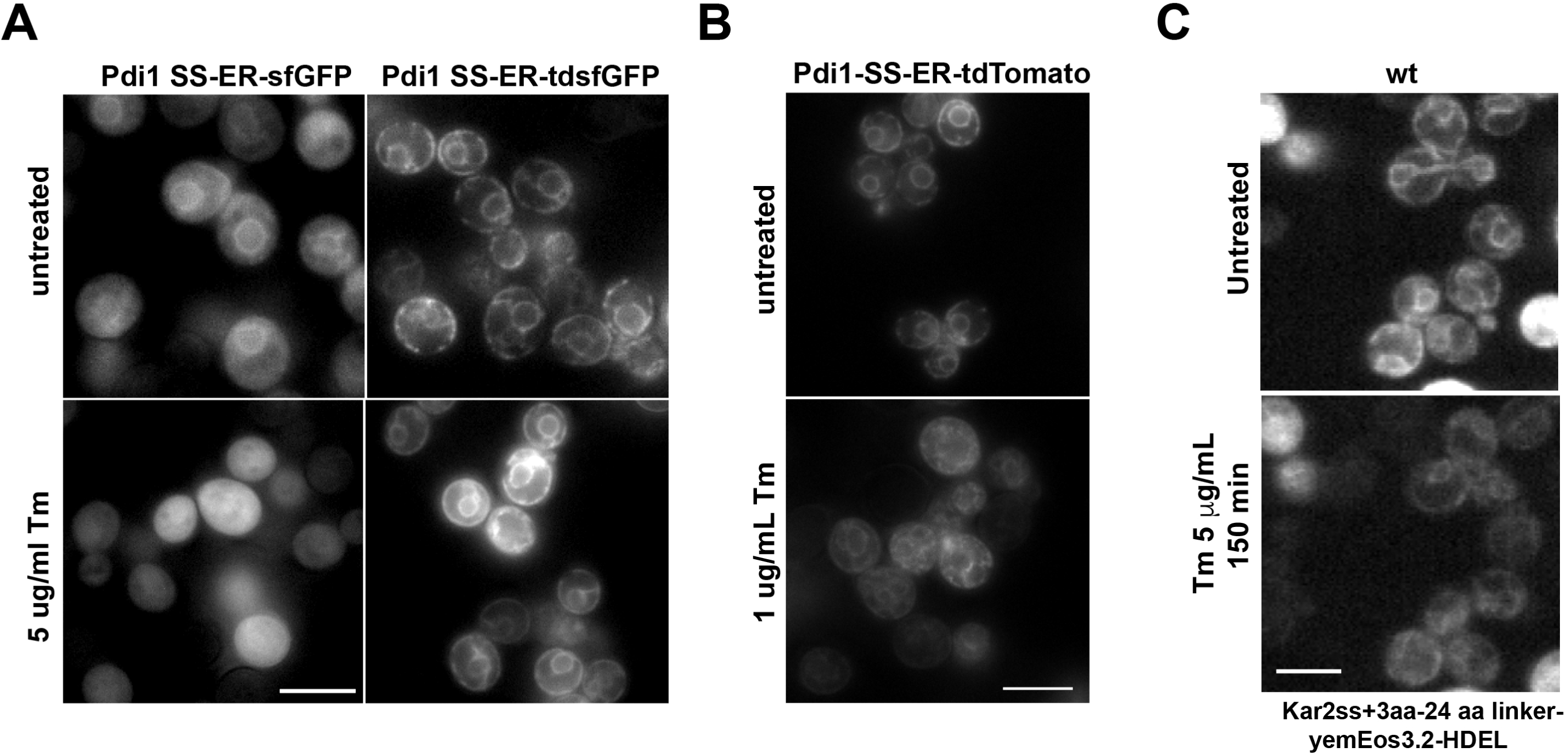
Tandem dimer FPs prevent stress stimulated cytosolic localization for a non-GFP FP and for the inefficient Pdi1 SS. (a) An FP construct with the poorly ER-localizing SS (Pdi1 SS-ER-sfGFP) exhibits dramatically improved ER localization at both steady state and during misfolded secretory protein accumulation (5 μg/mL Tm 2 h). (b) Maintenance of ER localization during ER stress is independent of the sfGFP sequence. The same strategy works with tdTomato (17), which has low amino acid sequence identity with sfGFP. (c) Adding the 24 a.a. GS12 linker protects photoconverted ER-localized mEos3.2 from relocating to the cytosol during Tm stress. Scale bars = 5 μm.

**Supplementary Table 1.**
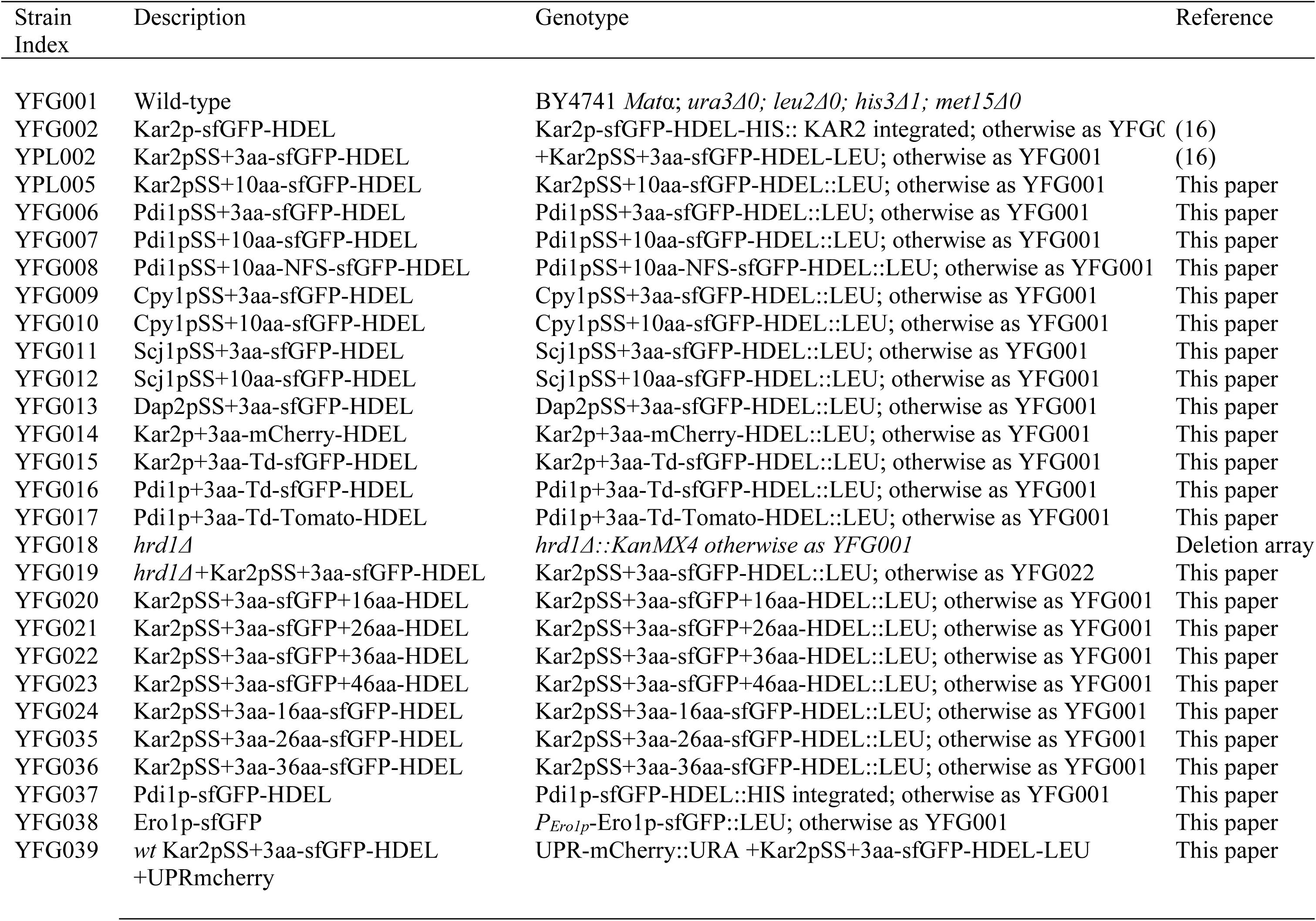
Yeast strains used in this study.

